# Expanding the toolbox for phycobiliprotein assembly: phycoerythrobilin biosynthesis in *Synechocystis*

**DOI:** 10.1101/2023.09.18.558311

**Authors:** Steffen Heck, Frederik Sommer, Susanne Zehner, Michael Schroda, Michelle M. Gehringer, Nicole Frankenberg-Dinkel

## Abstract

Phycobiliproteins (PBPs) play a vital role in light harvesting by cyanobacteria, which enables efficient utilization of photon energy for oxygenic photosynthesis. The PBPs carry phycobilins, open-chain tetrapyrrole chromophores derived from heme. The structure and chromophore composition of PBPs is dependent on the organism’s ecological niche. In cyanobacteria, these holo-proteins typically form large, macromolecular antenna complexes called phycobilisomes (PBSs). The PBS of *Synechocystis* sp. PCC 6803 (hereafter *Synechocystis*) consists of allophycocyanin (APC) and phycocyanin (PC), which exclusively harbor phycocyanobilin (PCB) as a chromophore. Investigations into heterologous PBP biosynthesis in *E. coli* have proven limiting with respect to PBP assembly and their functional characterization. Consequently, we wanted to engineer a platform for the investigation of heterologously produced PBPs, focusing on unusual, phycoerythrobilin (PEB)-containing light-harvesting proteins called phycoerythrins (PEs) in *Synechocystis*. As a first step, a gene encoding for the synthesis of the natural cyanobacterial chromophore, PEB, was introduced into *Synechocystis*. We provide spectroscopic evidence for heterologous PEB formation and show covalent attachment of PEB to the α-subunit of PC, CpcA, by HPLC and LC-MS/MS analyses. Fluorescence microscopy and PBS isolation demonstrate a cellular dispersal of PBPs with modified phycobilin content. However, these modifications have minor effects on physiological responses, as demonstrated by growth rates, oxygen evolution, nutrient accumulation, and PBP content analyses. As a result, *Synechocystis* demonstrates the capacity to efficiently manage PEB biosynthesis and therefore reflects a promising platform for both biochemical and physiological investigations of foreign and unusual PEs.

## Introduction

Cyanobacteria utilize light-harvesting complexes containing chlorophyll *a* as a chromophore, which absorb both blue (λ_max_=∼440 nm) and red light (λ_max_=∼678 nm). As cyanobacteria inhabit diverse ecological niches in terrestrial, marine, and freshwater environments, they need to adapt to the distinct light conditions in their habitats. To this end, they have developed specialized light-harvesting proteins known as phycobiliproteins (PBPs) (Apt et al., 1995). PBPs bridge the “green gap” between ∼500 nm and ∼650 nm, which is not efficiently absorbed by chlorophyll *a*, thereby enhancing light-harvesting efficiency (Glazer, 1977; Dagnino-Leone et al., 2022). In most cyanobacteria, these accessory pigments are organized via linker proteins in special light-harvesting antenna complexes, called phycobilisomes (PBSs). They funnel incoming light energy to the chlorophyll *a* containing photosystems’ reaction centers, thereby driving photosynthetic activity. The influence on photosynthesis depends on the size of the antenna complex as shown by studies with antenna truncation mutants (Page et al., 2012; Kirst et al., 2014). Variants of allophycocyanin (APC) form the PBS core, which is surrounded by up to six rods of phycocyanin (PC) and, in some cases, phycoerythrin (PE) (Sánchez-Baracaldo et al., 2019; Domínguez-Martín et al., 2022). PBPs are classified into different types based on the energy levels derived from their bilin chromophores (Sui, 2021). The first class is responsible for the absorption of high-energy light and consists of PEs (λ=540-570 nm) with covalently attached phycoerythrobilin (PEB) chromophores. Second are PCs (λ=610-620 nm), that are involved in harvesting intermediate-energy light and carry phycocyanobilin (PCB) as chromophores. And lastly, APCs (λ=650-670 nm) with covalently attached PCB, which enable the absorption of low-energy light (Peng et al., 2014). The incoming light is hierarchically processed from high-energy-to low-energy light within the PBS. Allophycocyanin B (AP-B) and the chromophorylated core-membrane linker L_cm_ (λ_max_=∼670 nm) serve as terminal emitters and pass the energy further to the chlorophyll *a* in the photosystems’ reaction centers (Ashby and Mullineaux, 1999). Most of the light absorbed by the PBS is channeled to photosystem II (PSII) (Glazer, 1984), which powers the hydrolysis of water to generate energy and producing oxygen as a byproduct (Shen, 2015).

The chromophores of the light-harvesting proteins represent a subtype of linear tetrapyrroles, known as phycobilins. The attachment of the chromophore to the apo-proteins requires specific ligation enzymes called PBP lyases. These PBP lyases ensure accurate attachment of the phycobilin to the corresponding cysteine residue with stereochemical precision (Scheer and Zhao, 2008). The phycobilins are derived from the cyclic tetrapyrrole heme which is cleaved by heme oxygenase yielding biliverdin IXα (BV) (Fig. 1) (Frankenberg-Dinkel, 2004). Ferredoxin-dependent bilin reductases (FDBRs) process BV to various phycobilin chromophores (Frankenberg et al., 2001; Kohchi et al., 2001). The PCB:ferredoxin oxidoreductase PcyA specifically converts BV to PCB (Frankenberg and Lagarias, 2003). The biosynthesis of PEB in cyanobacteria involves two distinct FDBR enzymes. The 15,16-dihydrobiliverdin:ferredoxin oxidoreductase, PebA, converts BV to 15,16-dihydrobiliverdin, which is then channeled to the PEB:ferredoxin oxidoreductase PebB, responsible for the formation of the final product, 3(*Z*) PEB (Dammeyer and Frankenberg-Dinkel, 2006). Contrary to the cyanobacterial two-step reaction, a single cyanophage encoded enzyme forms PEB via a phycoerythrobilin synthase (PebS) (Dammeyer et al., 2008).

**Figure 1:**
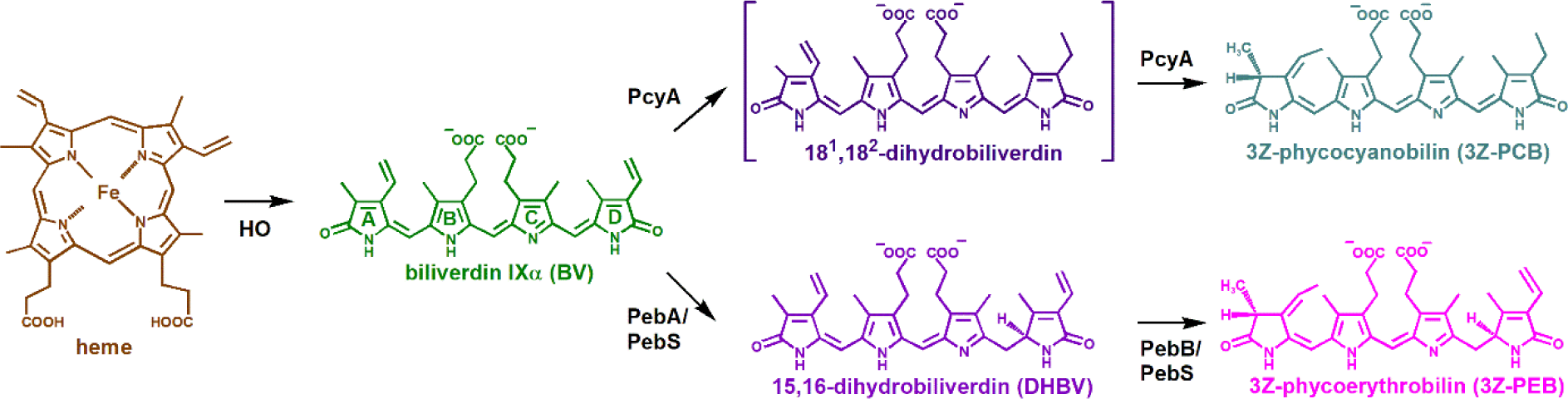
Biosynthesis of the linear tetrapyrroles PCB and PEB derived from heme. Cleavage of the cyclic tetrapyrrole heme by a heme oxygenase (HO) yields the linear tetrapyrrole BV, which is further converted to the phycobilins PCB and PEB by FDBR enzymes. In cyanobacteria, PCB is produced by one enzyme, PcyA, while PEB formation requires a two-step reaction catalyzed by PebA and PebB. In contrast, PEB catalysis is performed by a single FDBR from cyanophages called PebS.

Significant progress has been made in understanding the assembly of PBPs through *in vitro* studies using heterologous expression systems in *E. coli*. Those systems are widely employed to investigate the biochemical aspects of PBP biosynthesis and provide valuable insights into the assembly of light-harvesting proteins (Alvey et al., 2011; Biswas et al., 2011; Carrigee et al., 2021). However, we, and others, were repeatedly confronted with insolubility of proteins produced in these systems. Additionally, it has not yet been possible to assemble a complete holo-PBP (α− and β-subunit) in *E. coli*. In contrast, heterologous expression of foreign PBPs in cyanobacterial hosts offers an opportunity to investigate PBP assembly, their physiological roles *in vivo,* and allow for complementation of specific PBP mutant strains (Plank and Anderson, 1995; Puzorjov et al., 2021). These studies exclusively introduce PBPs that rely on the model organisms’ native phycobilins. However, a variety of PBPs depend on chromophore biosynthesis pathways not found in commonly used cyanobacterial model organisms like *Synechocystis* and *Synechococcus* sp. PCC 7002 (hereafter *Synechococcus*). Both strains only produce the phycobilin PCB and employ the PBPs PC and APC for light harvesting (Schluchter et al., 2010). Efforts have been made to modify endogenous phycobilin content by incorporating PEB biosynthesis pathways. For instance, in a marine *Synechococcus* strain, the introduction of *pebA* and *pebB* has detrimental effects on the pheno- and genotype (Alvey et al., 2011). On the other hand, a freshwater strain of *Synechocystis* lacking PC (ΔRod) and PCB biosynthesis was successfully engineered to exclusively produce PEB (Guo et al., 2022). In this case, the authors demonstrated attachment of PEB to the APC subunits ApcA and ApcB. Despite the significant progress already made, the ability to investigate heterologously expressed PEs both *in vitro* and *in vivo* within a single model system remains limited.

Consequently, we aim to develop a model system using *Synechocystis* wild type that enables the study of both the biochemical and physiological contribution of heterologously expressed PEs. In this study, we introduced PEB biosynthesis into *Synechocystis* via the *pebS* gene to expand the hosts phycobilin repertoire and present evidence for the formation of PEB. HPLC and LC-MS/MS analysis of purified PC thereby revealed covalent attachment of PEB to cysteine 84 of CpcA. The isolation of PBSs and co-localization studies via fluorescence microscopy showed that PBPs containing PEB mainly remained dispersed across the cytosol. To assess the effects of PEB production on cell physiology, we determined the growth rate and evaluated oxygen production, nutrient accumulation, as well as PBP content. While changes were observed in these aspects, the overall impact on cell physiology was minimal. These findings strengthen the potential of utilizing *Synechocystis* as a valuable platform for investigating heterologously produced PBPs containing PEB.

## Materials and methods

### Materials and reagents used

All chemicals used were reagent grade or better and purchased from Merck and Roth unless otherwise mentioned. Centrifugation steps were performed with a Hermle^TM^ Z32HK benchtop centrifuge unless otherwise mentioned. FPLC system and chromatography columns were acquired from Cytiva and HPLC columns from YMC Europe.

### Plasmid construction and conjugation

All Golden Gate cloning was performed utilizing the MoClo Plant Parts Kit (Kit #1000000044, addgene) and the CyanoGate Kit (Kit #1000000146, addgene). Transformed *E. coli* TOP10 cells were grown at 37 °C on LB medium supplemented with 1.5 % agar, spectinomycin (50 µg mL^−1^), carbenicillin (100 µg mL^−1^) or kanamycin (50 µg mL^−1^) as required. X-Gal (200 µg mL^−1^) was added for blue-white selection if needed. Liquid cultures of *E*. *coli* for plasmid extraction and conjugation were incubated at 37 °C and 160 rpm in LB medium under appropriate antibiotic selection. Golden Gate assembly reactions were performed according to Engler et al. (2014). To generate the level 0 coding sequence part, the *pebS* gene of the *Prochlorococcus* phage P-SSM2 was amplified by overlap extension PCR using Phusion^TM^ High-Fidelity DNA Polymerase. The amplicon was then assembled into the level 0 CDS acceptor vector pAGM1287 yielding pCG0.STH01. The native *cpc*-operon promoter (pC0.005) and terminator (pC0.078) were cloned up- and downstream of *pebS* (pCG0.STH01) into a level 1 position 1 (forward) acceptor vector pICH47732. Subsequently, the generated level 1 module was assembled in the level T acceptor vector pCAT.000 using the End-Link 1 in pICH41722, resulting in the cargo vector pCGT.STH02. Integrity of the new level 0 parts were verified by sequencing (Eurofins Genomics). Confirmation of the integrity of level 1 and level T constructs was performed by colony PCR and restriction enzyme digestion.

Genetic modification by conjugation into *Synechocystis* was performed according to Gale et al. (2019) with some minor modifications. The helper strain *E. coli* J53 [RP4] was grown in LB supplemented with 100 µg mL^−1^ ampicillin and 40 µg mL^−1^ kanamycin and the *E*. *coli* TOP10 cargo strains containing level T vectors were cultivated in LB with 50 µg mL^−1^ kanamycin. To generate a vector control strain (VC) of *Synechocystis*, the conjugative level T acceptor vector pCAT.000 was introduced. *Synechocystis* transconjugants were verified by colony PCR. All oligonucleotides used in this study are listed in Supplemental Data Table S1.

### Cultivation of cyanobacterial strains

*Synechocystis* liquid cultures were cultivated in BG-11 medium (Lea-Smith et al., 2016) at 30 °C under continuous warm-white LED light (40 µmol photons m^−2^ s^−1^) using an Infors Minitron incubator (Infors HT) with shaking at 120 rpm, unless otherwise specified. Transconjugated strains were grown in BG-11 medium supplemented with 50 µg mL^−1^ kanamycin. Maintenance of *Synechocystis* strains was performed on BG-11-agar (1.5 % w/v) with appropriate antibiotic selection under continuous white light (10 µmol photons m^−2^ s^−1^) at room temperature.

### Growth curves and chlorophyll *a* extraction

Growth experiments were performed in biological triplicates in 500 mL Erlenmeyer flasks. Initially, cultures were adjusted to an OD_730 nm_ of 0.1 (∼0.28 µg Chl *a* mL^−1^) in a culture volume of 100 mL. Growth of cultures was monitored for 7 days by measuring the absorbance at 730 nm. The Chl *a* extraction procedure was adapted from Zavřel et al. (2015). Samples were centrifuged at 6000 × g for 5 min at RT and the supernatant discarded. A 100 mg of 0.1 mm FastPrep silica beads (MPBIO, Germany) and 1 mL of 100% methanol were added to each sample pellet, followed by bead beating at 30 beats per second for 2 min (MM400 mixer mill, Retsch). After incubating overnight at 4 °C, the samples were vortexed and centrifuged at 15000 × g for 15 min. The UV-Vis absorption spectra of the cleared extracts, ranging from 250 nm to 800 nm, were measured (8453 UV-Visible system, Agilent Technologies). Chl *a* concentrations were calculated using α = 78.36 L g^−1^ cm^−1^ as the mass extinction coefficient in 100 % methanol (Li et al., 2012), after normalizing the samples to absorbance values at 780 nm.

### Measurement of cellular protein and glycogen content

For the quantification of cellular protein and glycogen content, the supernatant after cell lysis was used. The Bradford and anthrone assays were performed in 96-well plates following the procedure described by Herrmann and Gehringer (2019). Samples of the growth experiment were harvested by centrifugation at 6000 × g for 5 min at RT and the supernatant was discarded. A 100 mg of 0.1 mm FastPrep silica beads (MPBIO) and 1 mL of PBS buffer (154 mM NaCl, 5.6 mM Na_2_HPO_4_, 1.058 mM KH_2_PO_4_, pH 7.4) were added, and the samples lysed by bead beating as previously described. Subsequently, the samples were kept on ice in the dark for 1 h and then centrifuged at 15000 × g for 5 min at 4 °C. For protein determination, 50 µL of sample volume was mixed with 250 µL of Bradford reagent, incubated at RT for 20 min and the OD_595nm_ (Infinite F200 pro, Tecan) reading used to derive the protein concentration from a calibration curve (R^2^ = 0.99) of bovine serum albumin in lysis buffer. To quantify glycogen, 100 µL of soluble lysate was mixed with 250 µL of anthrone solution (2 mg mL^−1^ in 98 % sulfuric acid) and incubated at 80 °C for 30 min. After the reaction, optical densities were measured at 620 nm (FluoStar Omega, BMG Labtech). Glycogen concentrations were calculated from a calibration curve (R^2^ = 0.99) of glycogen from oyster in lysis buffer.

### Quantification of phycobiliproteins

The procedure of phycobiliprotein content determination was adapted from Zavřel et al. (2018) using fractions of the soluble lysate. The PC and APC contents were derived by UV-Vis spectroscopy (8453 UV-Visible system, Agilent Technologies) readings at 615 nm and at 652 nm, using the equations of Bennett and Bogorad (1973).

### Monitoring of oxygen release

Light response curves were generated by monitoring oxygen evolution of *Synechocystis* strains at RT under progressively increasing light intensities using the pulse amplitude modulation fluorometer Mini-PAM II (Heinz Walz GmbH). The dark-adapted cultures (20 min) were exposed to seven light intensities, which were incrementally increased over a period of 3 min to ensure stable signal acquisition. The cultures were stirred during the complete measurement. Triplicates were collected during mid-exponential growth phase (OD_730 nm_ = 0.4). Chlorophyll concentration was determined through Chl *a* extraction as previously described and the sample was adjusted to 2.5 µg mL^−1^ using fresh BG-11 medium (supplemented with 10 mM NaHCO_3_ and kanamycin if appropriate). A volume of 500 µL of the culture was transferred to a KS-2500 cuvette (Heinz Walz GmbH), with an optical oxygen sensor (OXR50-OI, PyroScience) connected. Following data acquisition, a moving average with a 17-point interval was calculated to compensate for minor oscillation during the assay. The oxygen evolution rate per hour was subsequently determined based on mg Chl *a*.

### Statistics and reproducibility

In physiological studies, the data points are presented as the mean values with standard deviation (± SD) derived from three independent biological replicates (n = 3). Evaluation of normal distribution was carried out using a Shapiro-Wilk test (Origin 2022, OriginLab). The analysis of variances was performed using an F-test, and the determination of significance was carried out using a two-tailed Student’s t-test (Excel 2016, Microsoft).

### Confocal fluorescence microscopy of living cells

Confocal fluorescence images were taken of exponentially grown cells, of which 5 µl of culture was spotted on an agarose pad (1 % low-melting agarose in BG-11), airdried and covered with a coverslip. All images were acquired using a Zeiss LSM880 AxioObserver confocal laser scanning microscope equipped with a Zeiss C-Apochromat 63x/1.4 oil-immersion objective. Samples were excited at 543 nm and phycobilin-derived emission for PEB was detected at 550-587 nm and PCB at 607-624 nm, while Chl *a* emission was recorded between 701-721 nm. Detection wavelengths for phycobilin-based fluorescence were optimized according to the sample’s emission spectra between 500-700 nm after excitation at 543 nm (Supplemental Data Figure S1). Acquisition settings for excitation and detection were identical for all images. Pictures were processed using the Zeiss software ZEN 2.3.

### Isolation of intact phycobilisomes

Preparation of intact PBS was performed as per Wallner et al. (2012). Cultures of *Synechocystis* WT and transconjugant strains (200 mL each) were harvested during late exponential growth phase (OD_730 nm_ = 0.7) by centrifugation at 6000 × g for 15 min at RT. The pellet was resuspended in 0.75 M potassium phosphate buffer (pH 7) and subjected to cell disruption through bead beating as described earlier. To remove cellular debris and silica beads, the sample was centrifuged twice at 15000 × g for 15 min at RT. Triton X-100 was added to the cleared lysates to a final concentration of 2 % (v/v) and the samples subsequently centrifuged as described above. Three mg protein of the PBS containing fractions were loaded onto a discontinuous sucrose gradient consisting of 1.5 M, 1 M, 0.75 M and 0.5 M sucrose in 0.75 M potassium phosphate buffer (pH 7). Ultracentrifugation was performed in a SW40 Ti swinging-bucket rotor at 130000 × g for 16 h at 20 °C (Optima L-80 XP, Beckman Coulter).

Fractions obtained from the sucrose gradient were characterized by UV-Vis spectroscopy, fluorescence emission spectra analysis, SDS-PAGE and Zn^2+^-Blot as described below.

### Purification of phycocyanin

The isolation of PC was performed using a modified version of the method described by Zhang and Chen (1999). Cells were harvested at an OD_730 nm_ of ∼1 and freeze-dried for storage. Freeze-dried samples were resuspended in 50 mM Na-phosphate buffer (10 ml/100 mg cells, pH 7) and subjected to sonication for 2 min with 9 cycles and 50 % power at 4 °C on ice (KE 76, Sonopuls HD2200, Bandelin). The lysate was the centrifuged at 16000 × g for 10 min at 4 °C. The supernatant was subjected to a two-step ammonium sulfate precipitation at 30 % and 70 % saturation. After each step samples were centrifuged at 50000 × g for 30 min at 4 °C. The pellet obtained from the 70 % saturation step was resuspended and dialyzed over night against 2 L of purification buffer (20 mM Bis-Tris buffer, pH 7). The dialyzed sample was loaded onto a HiTrap^TM^ DEAE Sepharose FF anion exchange column connected to a ÄktaPure FPLC system. After excessive washing with purification buffer, the samples were eluted with a stepwise increasing concentration gradient of NaCl in purification buffer (0 M, 0.1 M, 0.15 M, 0.2 M and 0.25 M). Fractions containing PC were applied onto a Superdex 200 increase 10/300 GL gel filtration column and samples were eluted with purification buffer at a flowrate of 0.2 ml min^−1^. Subsequently, purified PC was analyzed by SDS-PAGE, Zn^2+^ enhanced fluorescence, UV-Vis- and Fluorescence spectroscopy.

### SDS-PAGE and Zn^2+^ enhanced fluorescence

Proteins after PBS isolation and PC purification were separated by Bis-Tris SDS-PAGE in a 12 % separation gel to analyze the protein composition after PBS isolation and during PC purification. The proteins were either visualized by Coomassie Blue G-250 staining or transferred onto a PVDF-membrane for Zn^2+^ enhanced fluorescence (Berkelman and Lagarias, 1986). Therefore, the membrane was incubated in 1.3 M zinc acetate solution for 1 h and subsequently exposed to UV-light (312 nm) to visualize phycobilins covalently associated to proteins.

### Spectroscopic analyses

All absorbance spectra in a range of 250 nm to 750 nm were obtained using a UV-Vis spectrometer (Agilent Technologies, 8453 UV-Visible system). Fluorescence emission spectra between 555 nm and 750 nm were obtained using a fluorescence spectrometer (FP-8300, Jasco). For detection of PEB- and PCB-containing proteins, samples were excited with a wavelength of 550 nm. After data acquisition, values were normalized to the absorbance at 750 nm and mg protein of the fractions.

### Tryptic digest of PC and HPLC analysis of chromopeptides

Purified PC was precipitated in 80 % acetone (v/v) overnight at −20°C and centrifuged at 15000 × g for 15 min at 4 °C. Subsequently, the dried pellets were resuspended in 50 mM ammonium bicarbonate (pH 7.8). Digestion with trypsin from bovine pancreas (T4799, Sigma-Aldrich) was performed at 37 °C overnight (1:20 w/w trypsin to protein) in the dark. Trypsin was inactivated by adding 30 % (v/v) acetic acid and the resulting peptides were desalted using a Sep-Pak^®^ C18 3cc Vac cartridge (Waters Corporation) and lyophilized for HPLC analysis. Freeze-dried samples were resuspended in a solvent of 0.1 % formic acid in H_2_O. Peptides were applied onto a YMC-Triart C18 reverse phase column (10.0 x 2.1 mm) connected to a Shimadzu Prominence LC-20A HPLC system and eluted by a linear increasing gradient of Buffer B (0.1% formic acid in acetonitrile), from 15-60 % over 17 min, and from 60 % to 80 % over 5 mins, at a flowrate of 0.3 mL min^−1^. Elution of chromopeptides was followed by a built-in diode array detector, and 150 µl fractions were collected with a fraction collector. Prior to LC-MS/MS analysis, solvents were exchanged by evaporation and resuspension in a solvent of 1 % formic acid, 1 % acetonitrile and 0.1 % TFA in water.

### MS/MS analysis of PC chromopeptides

Mass spectrometry using an LC-MS/MS System (Eksigent 425 HPLC coupled to a TripleTOF 6600, AB Sciex, Darmstadt) was performed basically as described previously (Hammel et al., 2018). For peptide separation, a HPLC flow rate of 4 μl min^−1^ and 11 min gradients from 5% to 70% HPLC buffer B were employed (buffer A: 1% acetonitrile, 0.1% formic acid; buffer B: 90% acetonitrile, 0.1% formic acid) followed by washing and equilibration steps. MS1 spectra (350 *m*/*z* to 1500 *m*/*z*) were recorded for 250 ms and 20 MS/MS scans (100 *m*/*z* to 1500 *m*/*z*) were triggered in high sensitivity mode with a dwell time of 40 ms resulting in a total cycle time of 1100 ms. Analyzed precursors were excluded for 7 s, and singly charged precursors or precursors with a response below 300 cps were excluded completely from MS/MS analysis. PEB and PCB containing peptides were identified manually by filtering MS/MS spectra for PEB/PCB fragments and assigning MS1 precursor masses and MS/MS fragment masses to the peaks using PeakView2 software (AB Sciex, Darmstadt).

## Results

### Modified phycobilin biosynthesis impairs growth but promotes photosynthetic activity

In order to investigate the influence of foreign PEB biosynthesis in *Synechocystis*, the *pebS* gene from cyanophage P-SSM2 was introduced on a replicative plasmid under the control of the strong *cpc*-promoter (Vasudevan et al., 2019). The confirmed transconjugants already exhibited a slight difference in color, suggesting successful production of PEB. Consequently, the strain’s phototrophic growth was tracked over 7 days, through absorbance measurements at 730 nm and Chl *a* content. The time course profiles of the wild type (WT), vector control (VC) and *pebS* cultures are plotted below (Fig. 2A, B). The growth rates and doubling times during the mid-exponential phase were calculated from the Chl *a* content between 55 h and 30 h for the WT, 47 h and 23 h for the VC and 71 h and 47 h for the *pebS* strain (Fig. 2B) with statistical analyses conducted between the VC to the *pebS* growth rates.

**Figure 2:**
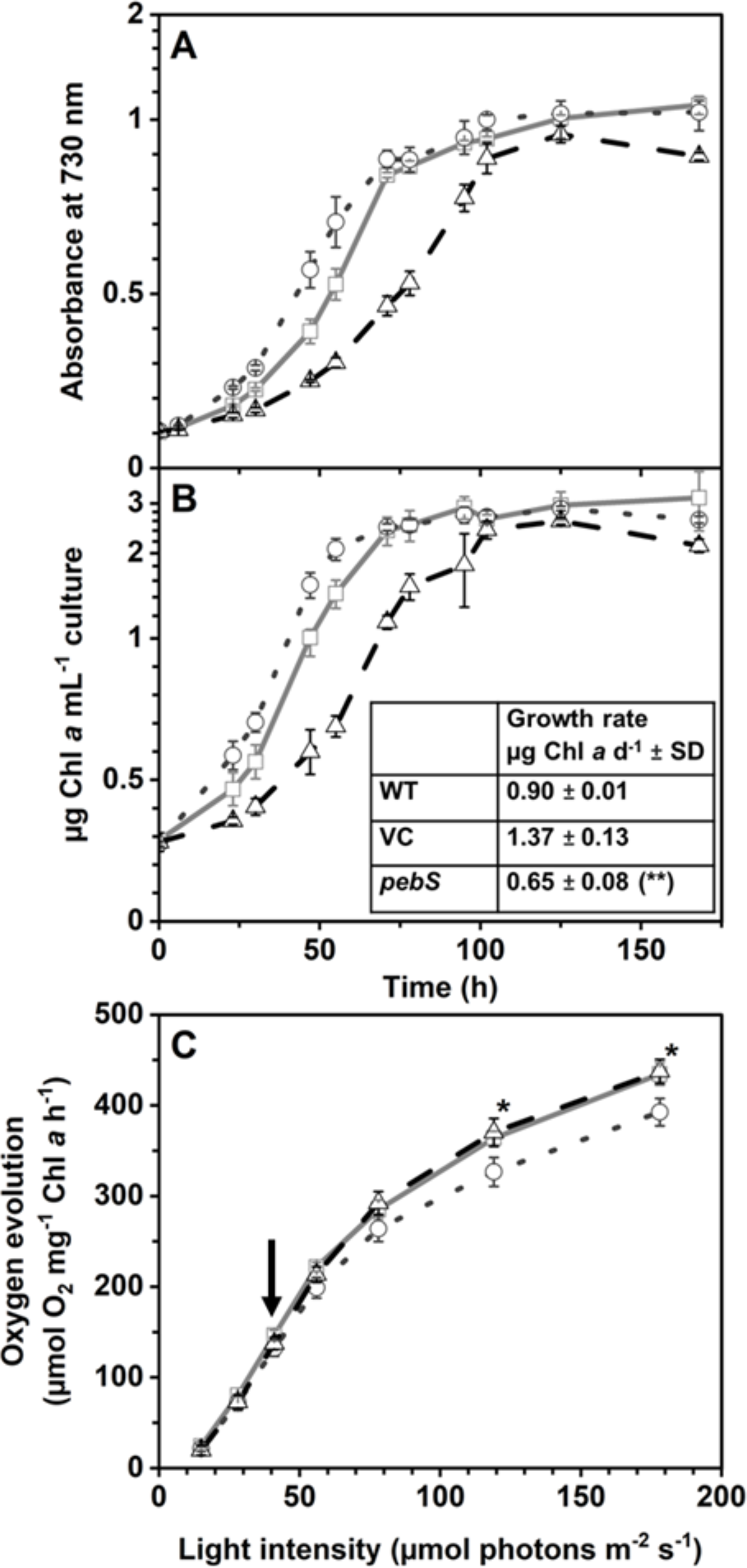
Phototrophic growth and light utilization of *Synechocystis* WT (grey line), VC (dotted line) and *pebS* strain (dashed line). Shown are mean values ± SD of *n* = 3. (**A**) Absorbance measurement at an optical density of 730 nm. (**B**) Chl *a* content per ml of culture. Growth rates are provided on the table inset (B). Statistical significance was conducted between the VC and the *pebS* strain. Asterisks indicate significant differences to the VC (** = p ≤ 0.01). (**C**) Oxygen evolution derived from water oxidation at PSII at incrementally increasing light intensities of mid-exponential phase cultures at ∼2.5 µg mL^−1^ Chl *a*. The arrow indicates O_2_ evolution at 40 µmol photons m^−2^ s^−1^, the culturing light intensity. Statistical significance was calculated between the VC and the *pebS* strain. Asterisks indicate significant differences to the VC (* = p ≤ 0.05).

The WT displayed a growth rate of 0.90 µg Chl *a* d^−1^ (± 0.01) with a doubling time of 0.77 d^−1^ (± 0.01). The VC exhibited the highest growth rate of 1.37 µg Chl *a* d^−1^ (± 0.13) and a doubling time of 0.51 d^−1^ (± 0.04). In contrast, the *pebS* strain demonstrated a growth rate of 0.65 µg Chl *a* d^−1^ (± 0.08) and a doubling time of 1.08 d^−1^ (± 0.14). The PEB producing strain showed no significant difference in growth rate, when compared to the WT, however, its growth rate was significantly lower (p ≤ 0.01) than that of the VC.

Based on changes in the growth rate we wanted to investigate the utilization of light energy during phototrophic growth (Fig. 2C). Therefore, the levels of oxygen release derived from water oxidation at PSII were assessed for all three strains over a range of light intensities. All three strains exhibited similar O_2_ release rates at the culturing light intensity (arrow, Fig. 2C). Once the light intensities rose above ∼50 µmol m^−2^ s^−1^, the VC steadily released less O_2_ into the medium compared to the WT and *pebS* strain. Significant differences became noticeable starting at a light intensity of 119 µmol photons m^−2^ s^−1^ (p ≤ 0.05). Considering the noticeable changes in growth rate but negligible variations in PSII activity, we sought to extend the evaluation of cellular viability by assessing accumulated nutrients and pigment content.

### Additional PEB biosynthesis influences pigment biosynthesis but not nutrient accumulation in *Synechocystis*

Cell viability is not only reflected by the growth rate, but also by the cellular reserves of nitrogen- and carbon-rich compounds. Therefore, the protein and glycogen content were determined from soluble cell lysates (Fig. 3A, B). Additionally, quantification of photosynthetically active pigments, such as PBPs, can provide insight into the physiological conditions during phototrophic growth. Therefore, we tracked the levels of the light-harvesting proteins PC and APC over time by UV-Vis spectroscopy (Fig. 3C, D). All values observed were normalized to µg Chl *a*.

**Figure 3:**
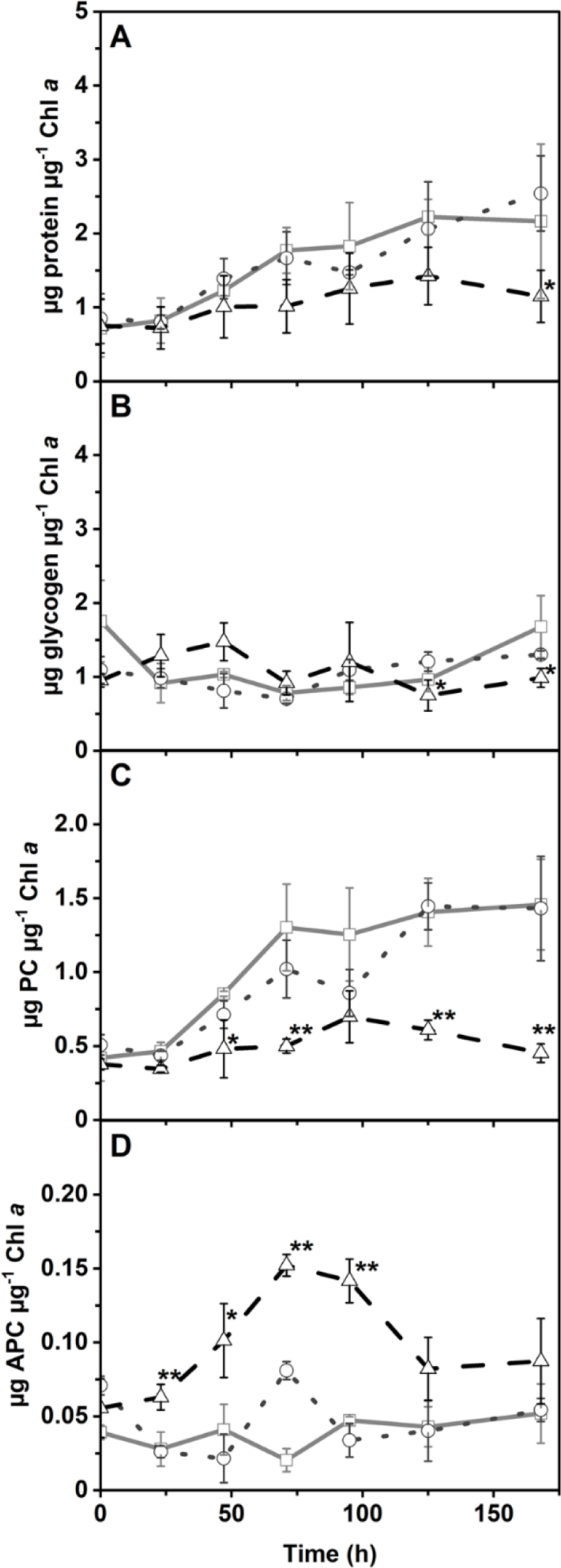
Assessment of parameters reflecting cell viability during phototrophic growth for Synechocystis WT (grey line), VC (dotted line) and the pebS strain (dashed line). Soluble cell lysates taken during the growth curve were used to determine the protein- (A), glycogen- (B), PC, (C) and APC- (D) content. Observed values were standardized to µg Chl a. Shown are mean values ± SD of n = 3. Asterisks indicate significant differences to the VC (* = p ≤ 0.05; ** = p ≤ 0,01).

Minor variations in protein content were observed between the VC (dotted line) and the *pebS* strain (dashed line) (Fig. 3A), with a significant difference recorded at 168 h (p ≤ 0.05). Similarly, the glycogen content (Fig. 3B) of the *pebS* strain was significantly lower than that of the VC at 125 h and 168 h (p ≤ 0.05). Overall, the *pebS* strain exhibited reduced levels of intracellular protein and glycogen as the cultures reached stationary phase when compared to VC and WT culture material.

Significant differences in PC content (Fig. 3C) were observed at 23 h post-inoculation (p ≤ 0.05), with the *pebS* strain exhibiting a reduction of ∼20 % in PC content compared to the VC. The difference in PC content persisted throughout the growth experiment, with reductions of 47 %, 41 %, and 32 % observed at 71 h, 125 h, and 168 h, respectively (p ≤ 0.01). In contrast, a two-fold increase in APC content (Fig. 3D) was observed in the *pebS* strain at 23 h (p ≤ 0.01). The amount of APC steadily increased, reaching a maximum value of 0.16 µg APC µg^−1^ Chl *a* at 71 h, thereby exceeding that of the VC by approximately 1.8-fold (p ≤ 0.01). In summary, the relative pigment contents, with respect to Chl *a*, were significantly altered in the *pebS* strain, independent of its growth phase.

### PEB is produced and induces PBP diffusion

The previous data provided evidence of a *pebS*-dependent effect on cell viability, including a notable reduction in PC levels, which suggested changes in phycobilin composition in the *pebS* strain. Therefore, we chose to investigate fractions of the soluble lysate by spectroscopic measurements to gather first-hand evidence of PEB production.

To reduce high background signals observed in whole cell UV-Vis analysis (data not shown), we focused our measurements on fractions of the soluble lysate to specifically detect PEB-related absorbance signals (Fig. 4A). After data acquisition, spectra were normalized to 750 nm and protein content. In the WT (black) and the VC (blue), the spectroscopic properties exhibited characteristic absorbance peaks at 622 nm and 652 nm, indicative of PC and APC, respectively. In contrast, measurements of the *pebS* strain lysate (red) revealed an additional signal at 562 nm besides the absorbance maxima at 622 nm and 652 nm. Furthermore, the absorbance per mg protein was reduced by approximately 60 % for PC at 622 nm and by around 50 % for APC at 652 nm in the PEB-producing strain.

**Figure 4:**
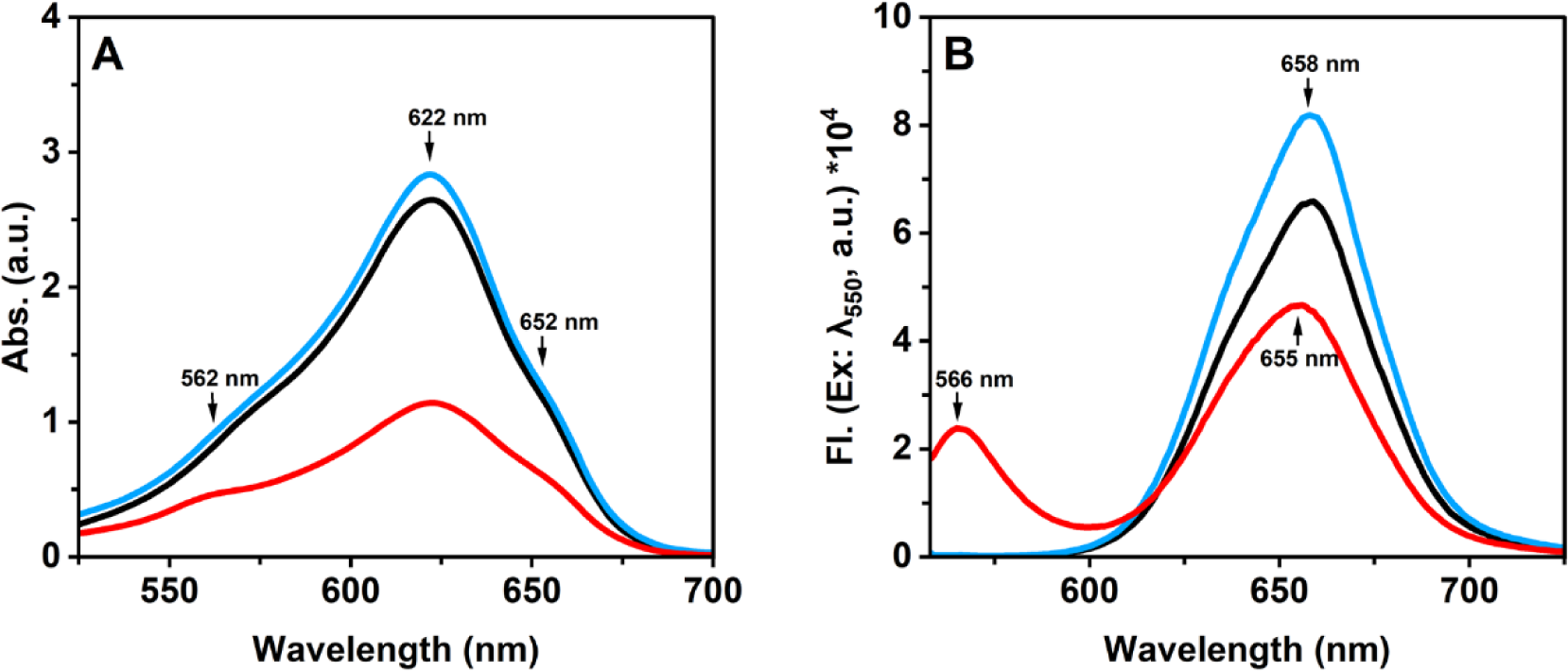
PEB production in *Synechocystis* WT and transconjugants. Soluble lysates of the WT (black), the VC (blue) and the *pebS* strain (red) were prepared from cultures in late-exponential phase for spectroscopic analyzes. Data was normalized to measurements at 750 nm and mg protein. **(A)** UV-Vis spectra of the soluble lysate in a range of 525 nm to 700 nm. **(B)** Fluorescence emission spectra of lysate fractions. The intensity of fluorescence emission between 558 nm and 725 nm after excitation at a wavelength of 550 nm is shown.

In contrast to free phycobilins, PBPs exhibit distinct fluorescent properties based on the type of phycobilin covalently attached to it. To investigate PEB-derived fluorescence emission, we performed fluorescence spectroscopy on the fractions used for UV-Vis measurements (Fig. 4B). In the spectra of the WT (black) and the VC (blue), a fluorescence peak maximum was observed at 658 nm, which indicates emission of APC. Interestingly, the measurements of the *pebS* strain (red) exhibited a shift in the emission peak from 658 nm to 655 nm. Additionally, a new emission signal at 566 nm was observed, indicating the expected signal of PEB-derived fluorescence.

As the initial spectroscopic investigations supported the synthesis and covalent linkage of PEB, we decided to observe the potential association of PEB to PBSs via confocal laser scanning microscopy (CLSM) (Fig. 5).

**Figure 5:**
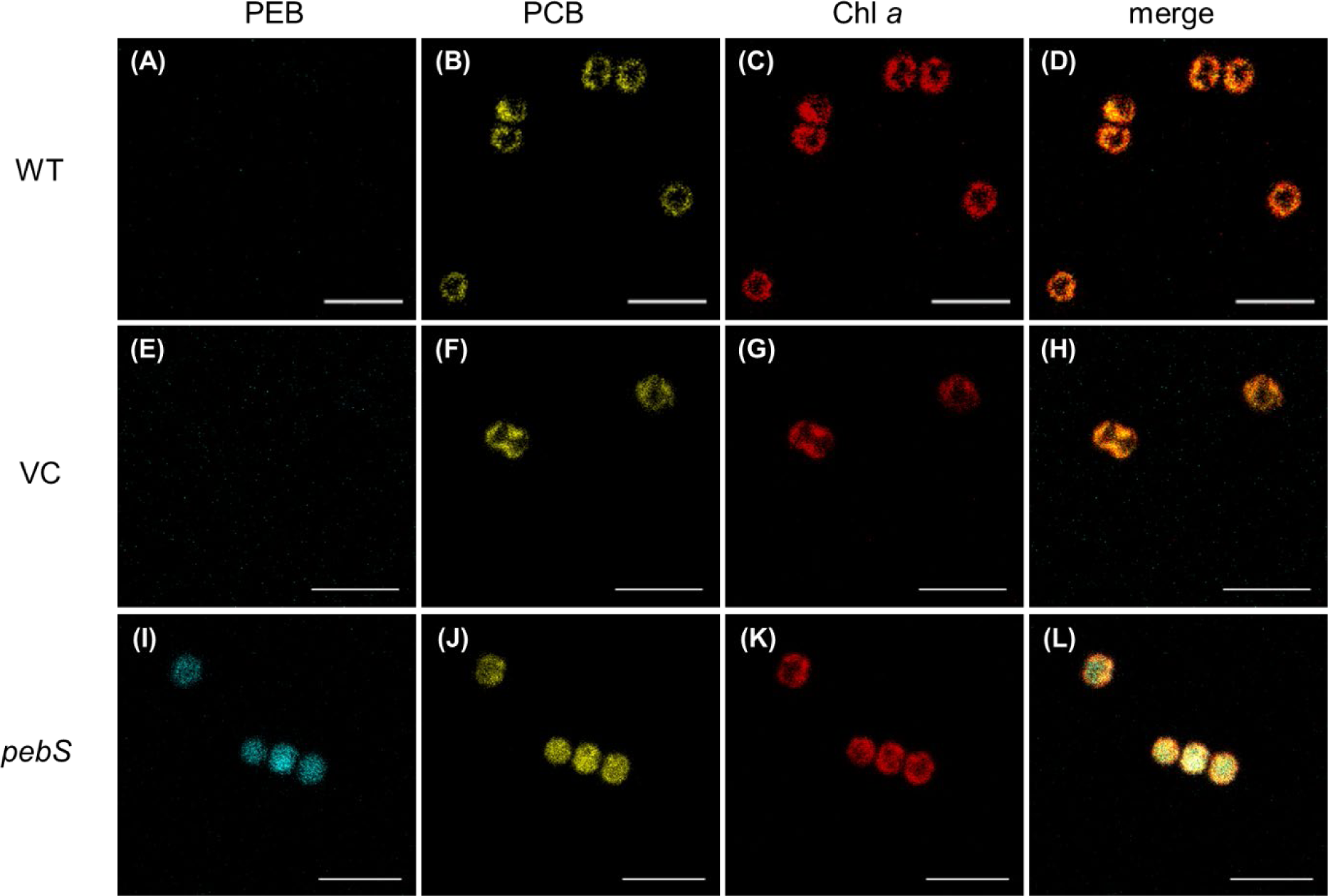
Confocal fluorescence images of *Synechocystis* WT (A to D), VC (E to H) and *pebS* (I to L). PBP-derived fluorescence signals were recorded for PEB between 550-587 nm (A, E, I), for PCB at 607-624 nm (B, F, J) and for Chl a at 701-721 nm (C, G, K) after excitation at 543 nm. Images were merged to visualize co-localization (D, H, L). Images are shown in false colors, showing fluorescence derived from PEB (blue), PCB (yellow) and chlorophyll A (red). Scale bars, 5 µm.

The CLSM profiles of *Synechocystis* WT and the VC revealed a distinct co-localization pattern of PCB-derived PBP fluorescence with Chl *a* fluorescence emission, indicating the association of PCB-containing PBSs with the thylakoid membrane (Fig 5D, H). While both control strains lacked PEB-based PBP emission signals (Fig 5A, E), the *pebS* transconjugant showed distinct signals distributed across the whole cell (Fig 5I). A similar pattern was observed for the PCB-derived PBP signals suggesting a highly reduced association to the thylakoids or even disassembly of the PBS. In contrast, the Chl *a* signal remained localized to the thylakoids (Fig 5K) as in in the control strains. Overlaying the images (Fig 5L) showed a highly reduced co-localization of PCB-derived PBP fluorescence and the thylakoid membrane in the PEB-producing strain. In contrast, PEB-derived PBP fluorescence signals were completely dispersed across the cell.

### PEB is absent in intact PBSs

As spectroscopic and CLSM fluorescence analyses clearly suggested the presence of PBP-bound PEB in the *pebS* transconjugant strain, we aimed to assess the integrity and chromophore composition of PBSs *in vitro* by isolating intact PBSs via sucrose density ultracentrifugation (Fig. 6).

**Figure 6:**
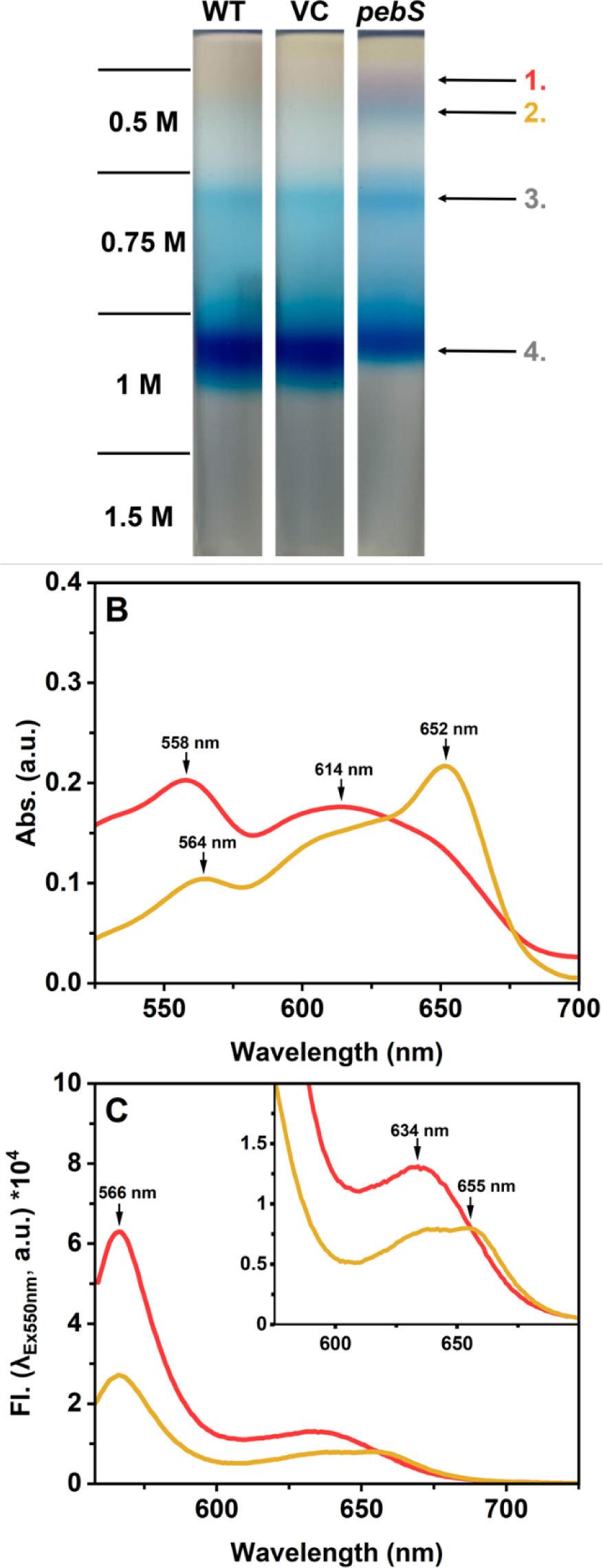
The influence of PEB biosynthesis on PBS profiles and its spectroscopic properties after ultracentrifugation. **(A)** Profiles of the PBS isolation from WT, VC and *pebS* strain samples after ultracentrifugation overnight in a discontinuous sucrose gradient (0.5 M, 0.75 M, 1 M and 1.5 M). Extracted fractions for downstream analyzes are indicated on the right. **(B)** UV-Vis spectra of fractions 1 (red) and 2 (orange) of the *pebS* strain in (A) between 525 nm and 725 nm. Data was normalized to absorbance at 750 nm and mg protein. **(C)** Fluorescence emission spectra of fractions 1 and 2 of the *pebS* strain in (A). Shown is the emission × 10^4^ between 558 nm and 725 nm after excitation at a wavelength of 550 nm. Recorded values were normalized to values at 750 nm and mg protein. Inset shows magnification of the region between 550 and 700 nm.

The obtained profiles of PBS complexes consisted of multiple pigment-based fractions, which were labeled as indicated in Fig 6A (4: 1 M; 3: 0.75 M, 2: 0.5 M bottom layer; 1: 0.5 M upper layer). The presence of intact PBS was consistently observed in all samples, represented by the blue color at a sucrose concentration of 1 M as already reported by Wallner et al. (2012). Signals observed at 0.75 M and 0.5 M sucrose concentrations indicated lower molecular mass assemblies of PBS components as demonstrated by Lea-Smith et al. (2014). Notably, new signals in magenta (Fraction 1) and blue color (Fraction 2) were detected in the 0.5 M sucrose concentration of the *pebS* strain sample. The fractions obtained after PBS isolation via ultracentrifugation were extracted from the corresponding sucrose layer and subjected to SDS-PAGE analysis to assess their protein composition, and subsequently transferred onto a PVDF membrane and treated with zinc-acetate to visualize phycobilins attached to peptides (Supplemental Data Figure S2 A).

Fractions 3 and 4 from all strains displayed no spectroscopic characteristics typical for the presence of PEB (Supplemental Data Figure S2 B-E). In contrast, the new fractions 1 and 2 in the 0.5 M sucrose layer of the PEB producing strain exhibited spectral properties characteristic of PEB. Fraction 1 (Fig. 6B, red) displayed an absorbance maximum at 614 nm, representing PC, with a small shoulder at 652 nm, indicative of APC. Additionally, a signal derived from PEB was observed at 558 nm. Similar spectroscopic features were observed in fraction 2 (in blue), where a distinct absorbance maximum at 652 nm was identified, with a shoulder at 614 nm. In contrast to the PEB-based signal at 558 nm in fraction 1, the peak was shifted to 564 nm in fraction 2 (Fig. 6B, orange). The subsequent fluorescence emission spectra (Fig. 6C) of the samples suggested covalent attachment of either PEB (566 nm) or PCB (634 nm and 655 nm) to the PBPs. Overall, spectral properties specific for PEB were only observed in the low-density assemblies consisting of PBPs in the *pebS* strain. Nevertheless, the presence of multiple spectroscopic signals derived from PEB suggested its covalent attachment to multiple PBP subunits. Due to the challenging nature in separating these two novel fractions, an alternative purification method was established to more clearly elucidate the site of covalent attachment of PEB to the PBP. To this end, PC was isolated employing ammonium sulphate fractionation in combination with ion exchange- and size exclusion chromatography.

### PEB binds to the α-subunit of PC at cysteine 84

In order to validate covalent linkage of PEB to PC, we isolated PC from the *pebS* strain and characterized it by SDS-PAGE, UV-Vis spectroscopy, and fluorescence emission measurements to assess purity and spectral properties. Subsequently, PC was digested, and tryptic peptides were further analyzed using HPLC and LC-MS/MS to identify PC-derived peptides containing covalently attached PEB chromophores.

The observed signals after SDS-PAGE at approximately 21 kDa and 18 kDa (Fig. 7A) correspond to the subunits CpcB and CpcA of PC, respectively. Both signals exhibited the characteristic fluorescence expected from Zn^2+^ enhanced fluorescence, confirming the presence of phycobilins covalently attached to PC. The UV-Vis spectrum of purified PC (Fig. 7B) exhibited two distinct absorbance maxima at 560 nm and 617 nm (red line), indicating the presence of PEB and PCB, respectively. Further evidence for covalent linkage of PEB is provided by the emission maximum at 566 nm in addition to a PCB-based signal at 639 nm in fluorescence analysis upon excitation at 550 nm (yellow line). Taken together, PC is capable of binding heterologously produced PEB despite the presence of the native chromophore PCB. Since phycobilins are typically attached to specific cysteine residues of the PBP subunits, we wanted to elucidate the binding sites of PEB to such conserved sites. Therefore, tryptic chromopeptides were separated by HPLC and the fractions were analyzed by LC-MS/MS (Supplemental Data Figure S3). Here, we observed one predominant peak exhibiting a PEB-derived absorbance signal. In MS analysis of this fraction (Supplemental Data Figure S3 C, D), we found the CpcA derived peptide C_84_AR with an *m/z* = 468,22^2+^ [M+2H]^2+^ (calculated *m/z* = 468,23^2+^ [M+2H]^2+^). Additionally, we found small proportions of the naturally occurring PCB-bound peptide of CpcA with the identical *m/z* = 468,22^2+^ [M+2H]^2+^ (Supplemental Data Figure S3 E, F). Overall, these data confirm the presence of two populations of α-subunits, the native ones with PCB bound, and those with attached PEB, suggesting a competition of both chromophores for the CpcA cysteine 84 binding site.

**Figure 7:**
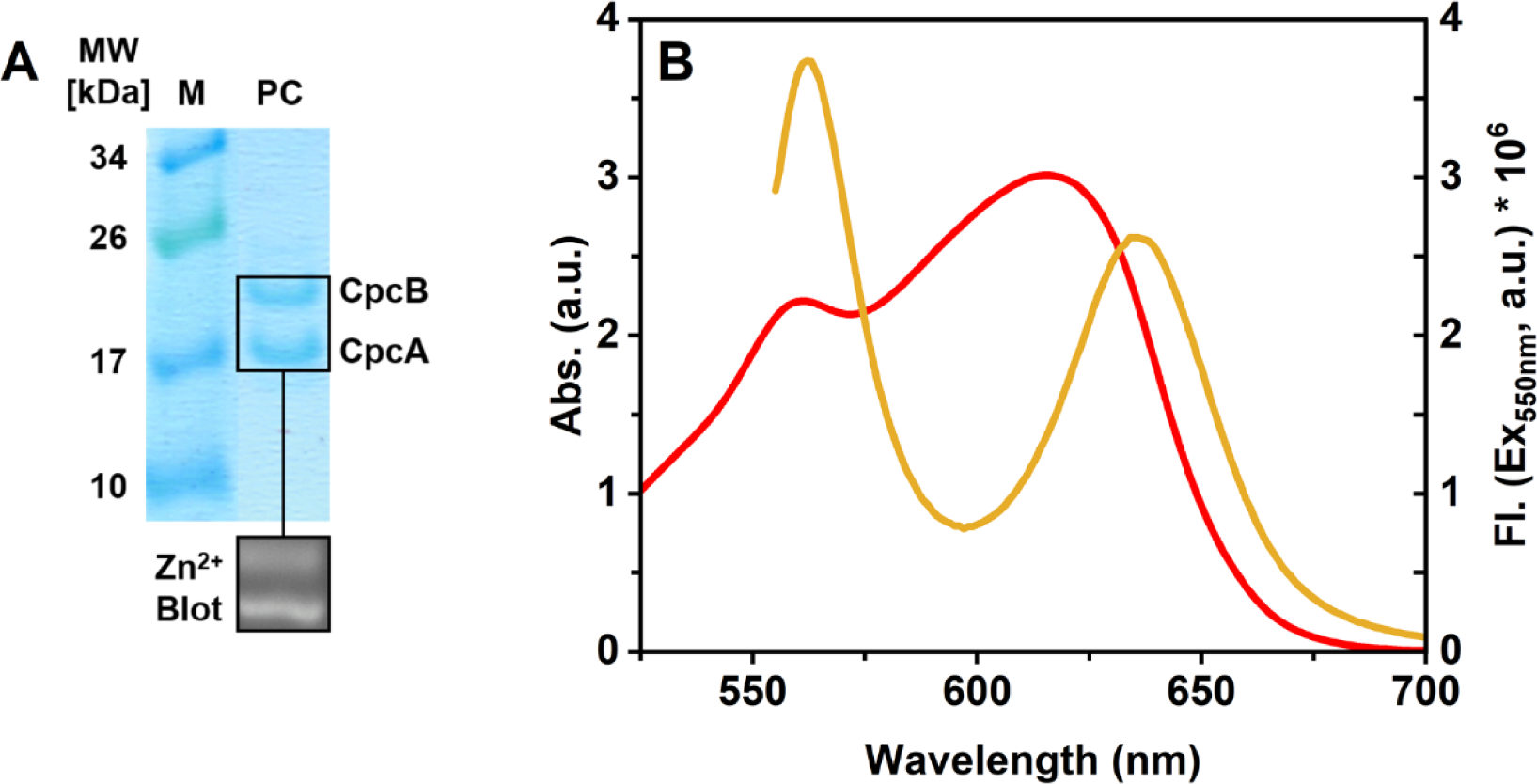
Modified PC from *Synechocystis pebS* strain after isolation. (**A**) SDS-PAGE analysis and Zn^2+^ enhanced fluorescence of purified PC after a three-step purification process. Line M indicates the molecular mass standard with corresponding MW in kDa. The signals in sample PC at ∼21 kDa and ∼18 kDa correspond to CpcB and CpcA, respectively. Covalent linkage of phycobilins to proteins was visualized via Zn^2+^ enhanced fluorescence after excitation at 312 nm. (**B**) UV-Vis spectroscopy and fluorescence emission measurements of purified PC. Data was normalized to mg protein. For UV-Vis analysis, spectral data of PC is shown between 525 nm and 700 nm (red). The fluorescence emission is displayed as relative intensity × 10^6^ between 558 nm and 725 nm after excitation at a wavelength of 550 nm (yellow).

## Discussion

Utilization of cyanobacterial model organisms such as *Synechocystis* for the investigation of heterologously expressed PBPs reflects a promising approach to investigate their physiological contribution to photosynthesis. This is especially important for PBPs originating from organisms that are not (yet) genetically accessible like *Prochlorococcus marinus*. The most frequently used platforms for studying these pigment-protein complexes utilize *E. coli*-based expression systems (Zhao et al., 2007; Wiethaus et al., 2010; Biswas et al., 2011; Kumarapperuma et al., 2022). Despite its great potential, this platform has still some limitations. Firstly, it is thus far not possible to produce a fully assembled PBP (α- and β-subunit) in *E. coli*, and secondly, this approach does not allow physiological characterization of PBPs and is therefore restricted to biochemical *in vitro* studies. The present study aimed at expanding the toolbox for PBP research by engineering *Synechocystis* to serve as a chassis for the investigation of light-harvesting proteins containing PEB as a chromophore. Besides the potential of utilizing the native cyanobacterial assembly machinery to enhance heterologous PBP biosynthesis, we sought to elucidate the physiological contribution of introduced PBPs to photosynthesis *in vivo*. As a first step to establish a cyanobacterial platform for the assembly of unusual PEB-containing PBPs, we introduced the *pebS* gene into *Synechocystis* to expand its phycobilin repertoire.

The production of PebS in *Synechocystis* leads to the formation of PEB (Fig. 4), which, in addition to the native chromophore PCB, is covalently bound to CpcA of PC (Supplemental Data Figure S3 C, D). These observations agree with previously reported studies that demonstrated covalent attachment of PEB to CpcA in *Synechococcus* (Alvey et al., 2011). Additionally, *in vitro* analyses based on heterologous expression systems in *E*. *coli* demonstrate very efficient attachment of PEB to CpcA (from *Synechocystis*) by the E/F-type lyases CpcE and CpcF from *Synechococcus* (Alvey et al., 2011). These findings, together with the data presented here, suggest a broad chromophore specificity of E/F-type lyases responsible for phycobilin attachment to Cysteine 84 of CpcA homologues. Here, the *Synechocystis* CpcE/F lyases are most likely responsible for the attachment of PEB to the cysteine residue 84 of CpcA (Fairchild and Glazer, 1994). Incorporation of PEB in CpcA resulted in impaired PBS assembly, which leads to free PEB-containing PBPs as shown by the dispersed PEB-derived signals in CLSM (Fig. 5) and PEB-containing low-density fractions after PBS isolation (Fig. 6). We hypothesize that binding of the heterologously produced PEB to PC influences the subunit structures and consequently prevents efficient PBS assembly. PEB and PCB are isomeric molecules, and they vary only in the length of their conjugated pi-electron system, ranging from the A-to the D-ring within the structure of the linear tetrapyrrole (Fig. 1). While the double bond between C15 and C16 in PCB makes the molecule more rigid, this linkage is reduced in PEB which gives the molecule increased flexibility (Tomazic et al., 2021). Consequently, this flexibility could lead to steric hindrance during the assembly of PBS if this structural change interferes with the protein-protein interface necessary for antenna formation. A similar case has already been reported for APC of a *Synechocystis* strain lacking PC (ΔRod) with attached PEB (Guo et al., 2022). APC naturally forms trimeric aggregates when PCB is bound, replacement with PEB inhibited their assembly beyond the monomer stage. In our study, where the non-native chromophore PEB was present in addition to the native PCB, we still observed a significant amount of native PBP that assembled into a PBS, although subjectively to a lower extent than in the WT (Fig. 6A). Overall, the total amount of PBP could be nearly identical to the WT situation (see also Fig. 3A), it is just not quantifiable with the methods employed here. Alternatively, introduction of PebS may lead to a competition for the substrate BV with the native PCB biosynthesis enzyme PcyA. This could result in reduced flux of BV into PCB biosynthesis and consequently a reduced synthesis of PC and APC holo-proteins necessary for PBS formation. Previous studies already suggest that PBP subunits lacking phycobilins are subjected to degradation based on an impaired subunit interaction (Plank and Anderson, 1995; Toole et al., 1998).

Although the integrity of the PBS is affected under modified phycobilin biosynthesis, its consequence for cellular physiology remains minor. The introduction of PEB biosynthesis slightly reduces the growth rate (Fig. 2B) but does not show a significant effect on cellular reserves of nitrogen (Fig. 3A), suggesting no influence on general biosynthesis of proteins. Additionally, the strain solidly performs oxygenic photosynthesis, as reflected by stable O_2_ release and illustrated by observed glycogen levels (Fig. 3B), which implies minimal interference in NADPH generation and CO_2_ fixation. This work is the first step towards heterologous PE assembly in a cyanobacterial model organism by introducing PEB biosynthesis in *Synechocystis* in addition to the host’s native PCB biosynthesis. It shows that *Synechocystis* can cope with non-native chromophore biosynthesis and presents a promising basis for introducing PE assembly components.

## Acknowledgements

The authors thank Prof. Dr. Annegret Wilde and Dr. Mai Watanabe (University of Freiburg) for providing the *E. coli* J53[RP4] helper strain for conjugation and Prof. Dr. Alistair McCormick (University of Edinburgh) for supplying *Synechocystis* sp. PCC6803.

## Author contributions

SH, MMG and NFD designed research, SH performed research, SZ conducted CLSM investigation and FS conducted mass spectrometry. MS provided tools and expertise; SH, MMG and NFD wrote the paper. All authors read and approved the final version of the manuscript.

## Data availability

MS data are available upon request from the authors.

